# Different processes shape prokaryotic and picoeukaryotic assemblages in the sunlit ocean microbiome

**DOI:** 10.1101/374298

**Authors:** Ramiro Logares, Ina M. Deutschmann, Caterina. R. Giner, Anders K. Krabberød, Thomas S. B. Schmidt, Laura Rubinat-Ripoll, Mireia Mestre, Guillem Salazar, Clara Ruiz-González, Marta Sebastián, Colomban de Vargas, Silvia G. Acinas, Carlos M. Duarte, Josep M. Gasol, Ramon Massana

## Abstract

The smallest members of the sunlit-ocean microbiome (prokaryotes and picoeukaryotes) participate in a plethora of ecosystem functions with planetary-scale effects. Understanding the processes determining the spatial turnover of this assemblage can help us better comprehend the links between microbiome species composition and ecosystem function. Ecological theory predicts that *selection*, *dispersal* and *drift* are main drivers of species distributions, yet, the relative quantitative importance of these ecological processes in structuring the surface-ocean microbiome is barely known. Here we quantified the role of selection, dispersal and drift in structuring surface-ocean prokaryotic and picoeukaryotic assemblages by using community DNA-sequence data collected during the global Malaspina expedition. We found that dispersal limitation was the dominant process structuring picoeukaryotic communities, while a balanced combination of dispersal limitation, selection and drift shaped prokaryotic counterparts. Subsequently, we determined the agents exerting abiotic selection as well as the spatial patterns emerging from the action of different ecological processes. We found that selection exerted via temperature had a strong influence on the structure of prokaryotic communities, particularly on species co-occurrences, a pattern not observed among communities of picoeukaryotes. Other measured abiotic variables had limited selective effects on microbiome structure. Picoeukaryotes presented a higher differentiation between neighbouring communities and a higher distance-decay when compared to prokaryotes, agreeing with their higher dispersal limitation. Finally, drift seemed to have a limited role in structuring the sunlit-ocean microbiome. The different predominance of ecological processes acting on particular subsets of the ocean microbiome suggests uneven responses to environmental change.

**SIGNIFICANCE STATEMENT:** The global ocean contains one of the largest microbiomes on Earth and changes on its structure can impact the functioning of the biosphere. Yet, we are far from understanding the mechanisms that structure the global ocean microbiome, that is, the relative importance of environmental *selection*, *dispersal* and random events (*drift*). We evaluated the role of these processes at the global scale, based on data derived from a circumglobal expedition and found that these ecological processes act differently on prokaryotes and picoeukaryotes, two of the main components of the ocean microbiome. Our work represents a significant contribution to understand the assembly of marine microbial communities, providing also insights on the links between ecological mechanisms, microbiome structure and ecosystem function.

## INTRODUCTION

The surface ocean microbiome is a pivotal underpinning of global biogeochemical cycles, thus being crucial for the functioning of the biosphere (1–4). The smallest ocean microbes, the picoplankton, have a key role in the global carbon cycle (4), being responsible for an important fraction of the total atmospheric carbon and nitrogen fixation in the ocean (5–7), which supports ~46% of the global primary productivity (8). Oceanic picoplankton plays a fundamental role in processing organic matter by recycling nutrients and carbon to support additional production as well as by channelling organic carbon to upper trophic levels through food webs (4, 5, 9).

The ocean picoplankton includes prokaryotes (both bacteria and archaea) and tiny unicellular eukaryotes (hereafter picoeukaryotes), which feature fundamental differences in terms of cellular structure, feeding habits, metabolic diversity, growth rates and behaviour (10, 11). Even though marine picoeukaryotes and prokaryotes are usually investigated separately, they are intimately connected through biogeochemical and food web networks (12–14). Overall, given the large effects picoeukaryotes can have on the populations of prokaryotes (and vice versa), it is highly relevant to determine whether or not their communities are structured by the action of similar ecological processes.

Ecological theory explains the structure of communities by a combination of four processes: *selection*, *dispersal*, *ecological drift* and *speciation* (15–17). Selection involves deterministic reproductive differences among individuals from different or the same species as a response to biotic or abiotic conditions. Selection can act in two opposite directions, it can constrain (homogeneous selection) or promote (heterogeneous selection) the divergence of communities (18). Dispersal, the movement of organisms across space, affects microbial assemblages by incorporating individuals originating from the regional species pool. Dispersal rates can be high (homogenising dispersal), moderate, or low [dispersal limitation] (18). Dispersal limitation occurs when species are absent from suitable habitats because potential colonizers are too far away (19). The importance of dispersal limitation increases as geographic scale increases (20). Ecological drift (hereafter *drift*) in a local community refers to random changes in species’ relative abundances derived from the stochastic processes of birth, death and offspring generation (15, 17). The action of drift in a *metacommunity*, that is, local communities that are connected via dispersal of multiple species (21), may lead to neutral dynamics (20), where random dispersal is the main mechanism of community assembly. In this neutral scenario, if dispersal is not limited, local communities will tend to resemble random subsamples of the metacommunity (20). Finally, speciation is the emergence of new species by evolution (15, 17), and it will not be considered hereafter as it is expected to have a small impact in the turnover of communities that are connected via dispersal (22).

The interplay of selection, dispersal and drift may generate different microbial assemblages that could feature diverse metabolisms and ecologies (16). The action of selection (in moderate to high strength) together with moderate rates of dispersal may generate a deterministic coupling between specific environmental conditions and combinations of species, a spatial pattern known as *species sorting* (23). In contrast, high or low levels of dispersal may produce the opposite effect, that is, a decoupling between abiotic environmental conditions (i.e. selection) and species assemblages. Particularly, high dispersal rates may maintain populations in habitats to which they are maladapted (21). Inversely, low dispersal rates may preclude species from reaching suitable habitats, leading to species assemblages that become more different as the geographic distance between them increases (distance decay). Still, both selection and dispersal limitation can generate distance decay (24). Drift is expected to cause important random effects in local community composition in cases where selection is weak and populations are small (16, 25).

Whereas global-ocean connectivity patterns reveal the importance of dispersal in structuring communities in the upper ocean (26), our understanding of the relative role of selection, dispersal and drift in structuring the global-ocean microbiome is still poor (24, 27, 28). Multiple studies in diverse environments indicate that selection has a major role in structuring prokaryotic communities (23, 24), although there is also evidence pointing to drift as having a structuring role (29, 30). Here, we examine the mechanisms shaping the sunlit global-ocean microbiome by addressing the following questions: What are the relative roles of selection, dispersal and drift in shaping assemblages of prokaryotes and picoeukaryotes? What environmental variables exert selection? What spatial patterns emerge from the action of these three processes? We hypothesize that the major organismal differences between picoeukaryotes and prokaryotes (11) should result in a different relative importance of selection, dispersal and drift in structuring their communities. Specifically, we hypothesize that the lower capacity for dormancy of picoeukaryotes (11, 31) should result in a larger dispersal limitation when compared to prokaryotes, given that dormancy may allow prokaryotes transiting through large geographic areas that are unsuitable for growth. Furthermore, given that some prokaryotes may engage into metabolic cooperation (32), an uncommon behaviour in picoeukaryotes, we expect selection to generate more co-occurrences among prokaryotes than among picoeukaryotes.

## RESULTS

### Processes shaping the surface global-ocean microbiome

We analysed the prokaryotes and picoeukaryotes composing this microbiome in 120 tropical and subtropical stations sampled during the Malaspina-2010 expedition (33) **[Fig. S1]** by *Illumina* High-Throughput sequencing of 16S and 18S rRNA-genes. Based on these genes, we delineated Operational Taxonomic Units (OTUs) as species proxies (see Methods). We applied an innovative methodology based on phylogenetic and species turnover (22) that allowed us to quantify the relative importance of selection, dispersal and drift (See Methods).

We found that selection, dispersal and drift played a similar role in structuring prokaryotic communities, while dispersal limitation was the dominant force structuring picoeukaryotic communities **(Fig. 1)**. Selection explained ~34% of the turnover of prokaryotic communities, and ~17% of that in picoeukaryotes **(Fig. 1)**. Heterogeneous selection had a relatively higher importance in structuring picoeukaryotes as compared to prokaryotes (~16% vs. ~9%, respectively), while homogeneous selection was more important in structuring prokaryotic (~24%) than picoeukaryotic (~1%) communities **(Fig. 1)**. Dispersal limitation was by far the most important process structuring picoeukaryotic communities (~76%), while this process had a lower importance in prokaryotes (~35%) **[Fig. 1]**. Drift explained a larger fraction of community turnover in prokaryotes (31%) than in picoeukaryotes (~6%) **[Fig. 1]**. Homogenizing dispersal had a very limited role in the structuring of the global ocean microbiome (<1% for both picoeukaryotes and prokaryotes).

**Figure 1.**
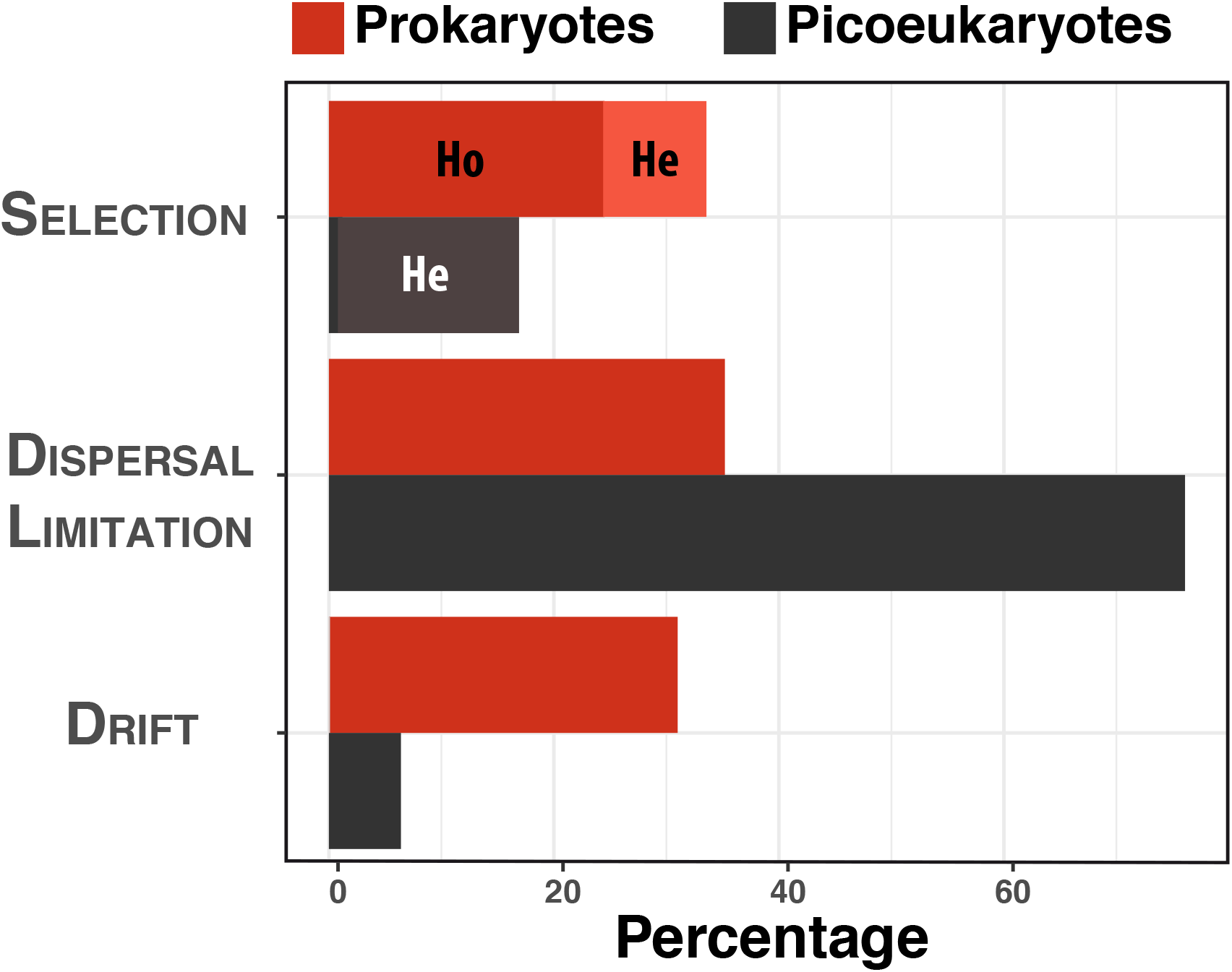
Relative importance of the processes shaping the tropical and subtropical sunlit-ocean microbiome. Percentage of the spatial turnover governed by different processes in prokaryotes and picoeukaryotes. Ho = Homogeneous selection, He = Heterogeneous selection (light red and light black).

Given that communities of prokaryotes and picoeukaryotes were predominantly structured by different processes, we expected both groups to present contrasting ß-diversity patterns. Accordingly, we found only moderate correlations between Bray-Curtis and generalized UniFrac (gUniFrac) ß-diversity indices between picoeukaryotes and prokaryotes (Bray Curtis: *ρ*=0.58, gUniFrac: *ρ* =0.61, p=0.01, Mantel tests; **Fig. S2**). Rare species tend to occupy less sites than more abundant species (34), therefore communities featuring different proportions of abundant and rare species may display different spatial turnover. We found that picoeukaryotes had proportionally more regionally rare species (here defined as those with mean abundances below 0.001%) than prokaryotes (71% vs. 48% of the species respectively) **[Table S1, Fig. S3]**. This is consistent with the observation that picoeukaryotes had more restricted species distributions (i.e., occurring in <20% of the communities) than prokaryotes [95% vs. 88% of the species respectively] (**Table S2; Fig. S3**).

### Selection acting on the microbiome

We investigated the abiotic agents exerting selection in the ocean microbiome by analysing the compositional differences between communities (ß-diversity) together with a set of environmental variables considered in the *Meta-119* dataset (See Supplementary Information). We used different ß-diversity indices (Bray-Curtis, TINA_w_, PINA_w_, gUniFrac; See Methods), as each captures distinct features of community differentiation. Water temperature was the most important driver of selection on prokaryotes **(Fig. 2)**. Furthermore, water temperature appeared to affect prokaryotic associations, given that the association-aware ß-diversity index TINA_w_ (35) explained ~50% of community variance (PERMANOVA *R^2^*) **[Fig. 2]**, while other ß-diversity indices tested that do not consider species associations explained considerably lower proportions **(Fig. 2)**. In contrast, temperature had limited effects on picoeukaryotic species associations **(Fig. 2)**. Our results were further confirmed by independent data from the global surface-ocean collected during the TARA Oceans expedition (36), as both the Malaspina and TARA Oceans datasets displayed stronger positive correlations between TINA_w_ and water-temperature differences in prokaryotes than in picoeukaryotes **[Fig. 3]**. This indicates that locations with similar temperatures feature co-occurring prokaryotic species, with this pattern disappearing as the temperature difference between stations increases.

**Figure 2.**
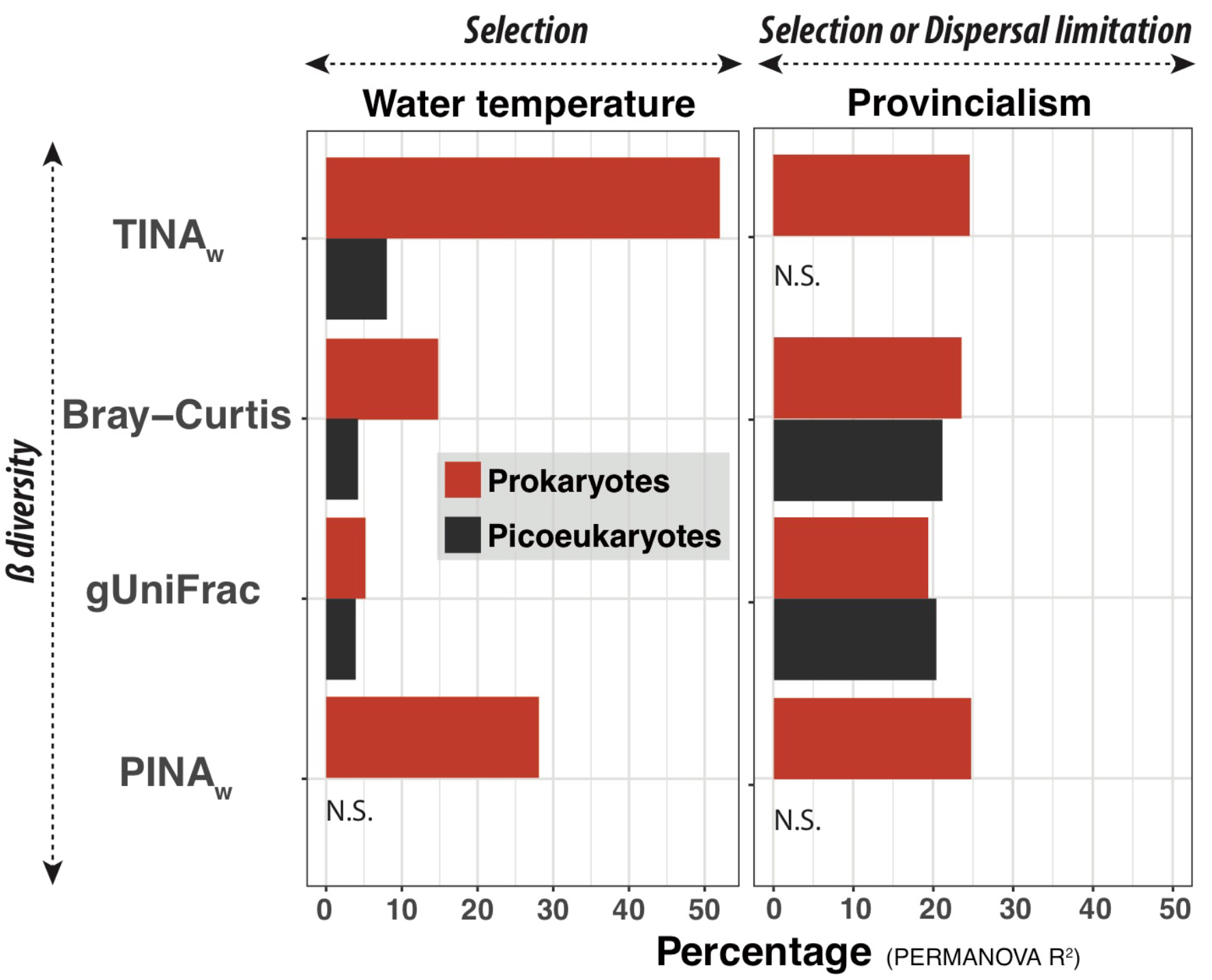
Water Temperature and Provincialism were the most important variables explaining the structure of the tropical and subtropical surface-ocean microbiome. Percentage of variance in picoeukaryotic and prokaryotic community composition (PERMANOVA R^2^) explained by Water Temperature and Longhurst Provinces when using different β-diversity metrics. Figure based on the *Meta-119* dataset (see Methods). TINA_w_: TINA weighted, gUniFrac: Generalized Unifrac, PINAw: PINA weighted. N.S. = Non-Significant. Note that TINA_w_ captures a significantly higher proportion of community variance than Bray-Curtis in prokaryotes.

**Figure 3.**
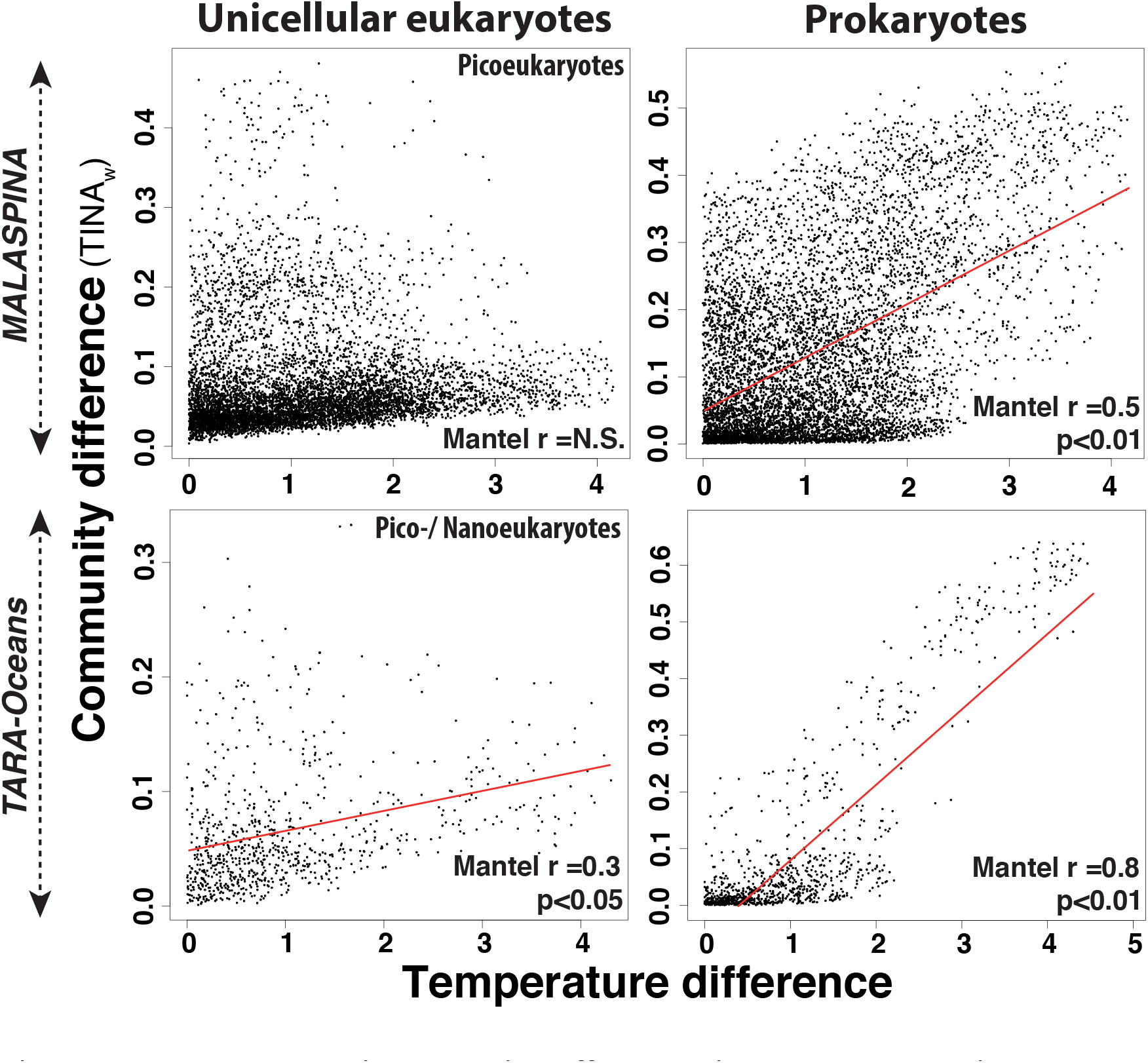
Temperature-driven selection affects species co-occurrences in prokaryotes but not in small unicellular eukaryotes. Community differences (Tina-weighted dissimilarities) vs. temperature differences (Euclidean distances) for both small unicellular eukaryotes and prokaryotes sampled during the Malaspina and TARA Oceans expeditions. NB: While only picoeukaryotes were contemplated in Malaspina (cell sizes <3 μm), TARA Oceans data included pico- and nanoeukaryotes (cell sizes <5 μm). Pico- and nanoeukaryotes from both expeditions (left panels) displayed low or no correlations between TINA_w_ distances and temperature differences (Note Mantel test results in the figures). On the contrary, prokaryotes (right panels) had high to moderate correlations between TINA_w_ distances and temperature differences. The regression line is shown in red (Malaspina microbial eukaryotes N.S., Malaspina Prokaryotes R^2^=0.3, TARA microbial eukaryotes R^2^=0.1, TARA Prokaryotes R^2^=0.7; p<0.05).

Further analyses, exploring 17 environmental variables from 57 stations (Supplementary Information**, Fig. S4)**, showed that fluorescence (a proxy for Chlorophyll *a* concentration) explained 31% of PINA_w_-based prokaryotic community variance (PERMANOVA *R^2^*), while it was not significant for picoeukaryotes **(Fig. S5**; PINA_w_ is a phylogeny-based ß-diversity index, See Methods). The remaining combinations of environmental variables and ß-diversity metrics explained a minor fraction of community variance, suggesting that abiotic selection, at the whole microbiome level, operates via a very limited set of environmental variables, largely temperature.

The finding that selection via temperature influences species associations particularly in prokaryotes suggests that prokaryotes and picoeukaryotes may show different patterns of species co-occurrences and co-exclusions in association networks (37). We found that prokaryotes were more associated between themselves than picoeukaryotes in networks considering co-occurrences and co-exclusions as well as in networks including only co-occurrences **(Fig. S6)**. Specifically, in networks including both co-occurrences and co-exclusions, prokaryotes featured ~33% of connected species (i.e. prokaryotic species with at least one association to another prokaryotic species) and an average number of associations per species (i.e. average degree) of ~14, while picoeukaryotes displayed ~17% of connected species and an average degree of ~8 **(Table S3**; **Fig. S6)**. Networks including co-occurrences only displayed similar patterns **(Table S3**; **Fig. S6)**. The prokaryotic network was more modular, containing a higher number of highly-connected clusters of species than the picoeukaryotic network **(Table S3)**.

The previous results were supported by analyses using the Maximal Information Coefficient [MIC], which quantifies a wide array of functional and non-functional, linear and non-linear, associations (38). MIC results indicated that prokaryotes had more associations between themselves than picoeukaryotes **(Table S4)**, a pattern that was also observed in other data from the upper global ocean collected during the TARA Ocean expedition **(Table S5)**. Most associations detected by MIC were non-linear [defining nonlinearity as MIC-ρ^2^ >0.2; (38)] **[Table S4** & **S5]**, pointing to complex associations that may be missed by classic correlation analyses, which evaluate linear relationships.

### Selection acting on species

The potential effects of selection on single species was evaluated by determining their individual correlations with multiple environmental variables using the MIC (38). In these analyses, temperature was the variable with the highest number of associated prokaryotic species (1.7%) when considering a MIC threshold ≥0.4, representing ~17% of the total estimated species abundance, while picoeukaryotic species displayed a considerable smaller proportion (~0.3% of the species representing ~5% of the estimated species abundance) [**Fig. S7]**. Picoeukaryotic and prokaryotic species also had associations with oxygen, conductivity and salinity **(Fig. S7)**, which co-vary with temperature. The remaining environmental variables tested had limited associations with individual species **(Fig. S7)**, thus agreeing with our previous results suggesting that selection on the surface ocean microbiome operates via a limited set of environmental variables, with a dominant role for temperature. Prokaryotes featured proportionally more individual-species associations with environmental parameters than picoeukaryotes **(Fig. S7**), thus pointing to a stronger abiotic selection pressure on prokaryotes than on picoeukaryotes in the surface global ocean. Our results were further validated by analyses using data from the global TARA Oceans cruise, which indicated that prokaryotic species were associated predominantly with temperature and oxygen, while small unicellular eukaryotes had limited associations to multiple variables (Temperature, Salinity, Oxygen, Nitrate & Chlorophyll; **Table S6**).

### Dispersal limitation

Our quantifications indicated that dispersal limitation was almost twice as important in structuring picoeukaryotic than prokaryotic communities **(Fig. 1)**. Environmental conditions between pairs of adjacent stations over the trajectory of the cruise, typically separated by 250-500 km, are generally comparable (i.e. selective differences between stations tend to be low in the tropical and subtropical ocean). Therefore, compositional differences between neighbouring communities could manifest dispersal limitation. Following these premises, we analysed the change in picoeukaryotic and prokaryotic community composition along the trajectory of the cruise by comparing each community to the one sampled immediately before in a sequential manner (i.e. sequential β-diversity) **[Fig. 4]**. Both picoeukaryotic and prokaryotic communities displayed variable amounts of sequential β-diversity **(Fig. 4, Panels A and B)**, although picoeukaryotes featured, on average, a higher sequential β-diversity than prokaryotes **(Fig. 4, Panel C)**. This is concordant with the overall mean β-diversity, which was significantly higher for picoeukaryotes than for prokaryotes **(Fig. S8)**.

**Figure 4.**
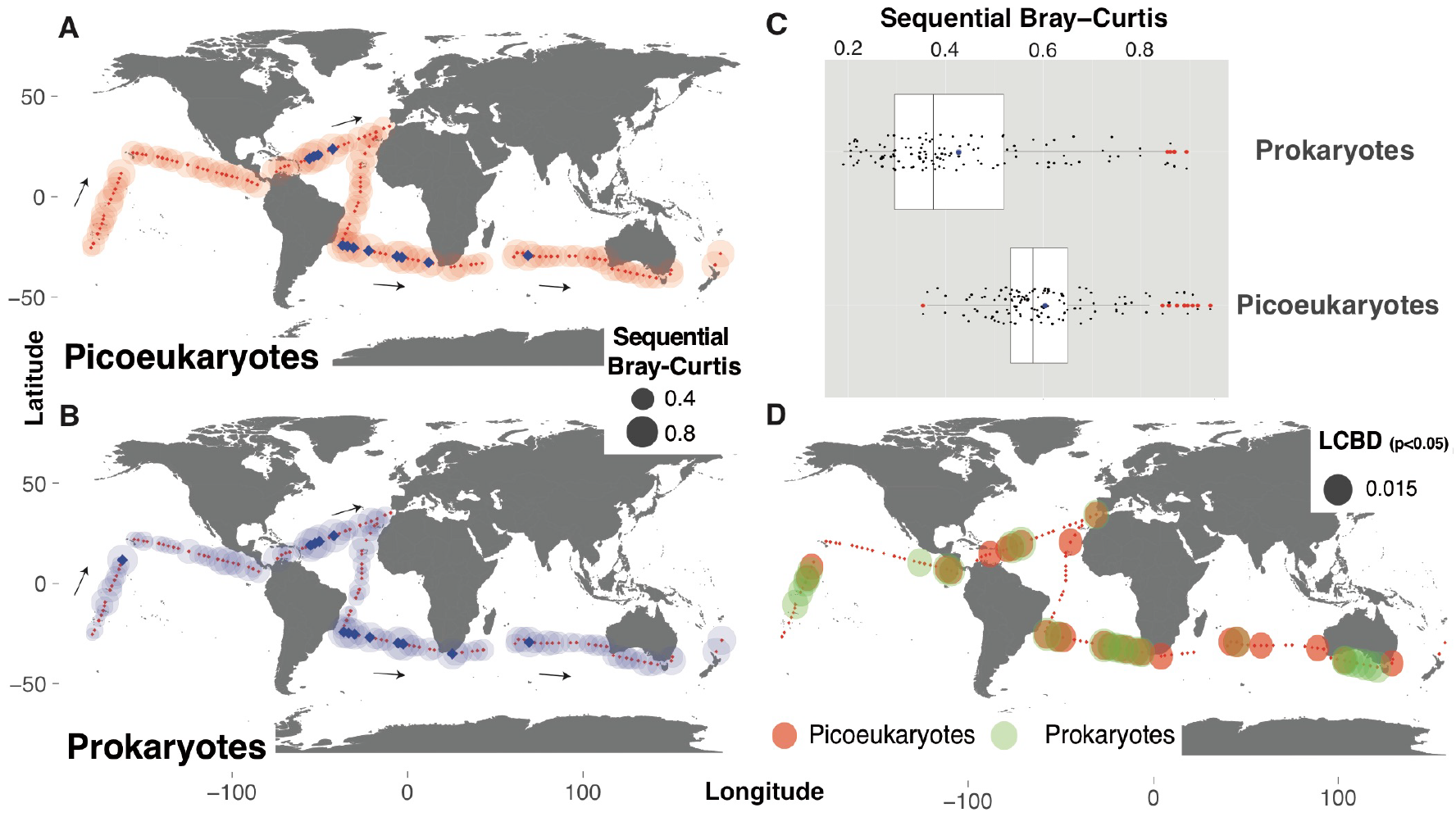
Spatial patterns likely manifesting differential dispersal or strong selective change. Panels A-C: Sequential change in community composition across space (sequential β-diversity). Communities were sampled along the R/V Hespérides trajectory (Panels A and B, black arrows), and the composition of each community was compared against its immediate predecessor. In Panels A and B, the size of each bubble represents the Bray-Curtis dissimilarity between a given community and the community sampled previously. Blue squares in Panels A and B represent the stations where pairwise β-diversity displayed abrupt changes (Bray Curtis values >0.8 for picoeukaryotes and >0.7 for prokaryotes). Abrupt changes coincided in a total of 11 out of 14 stations for both picoeukaryotes and prokaryotes, while one station presented marked changes for picoeukaryotes and two for prokaryotes. Panel C summarizes the sequential Bray-Curtis values for prokaryotes and picoeukaryotes [Means were significantly different between domains (Wilcoxon text, p<0.05)]. Panel D shows the 36 stations featuring a comparatively large contribution to the overall β-diversity (Local Contribution to Beta Diversity (40); p<0.05).

### Simultaneous action of selection and dispersal limitation

When geographic distance is correlated with environmental variation, spatial community variance may be the manifestation of both selection and dispersal limitation. We analysed community variance associated to different marine biogeographic provinces, as defined by Longhurst (39) based on nutrient concentration, structure of the water column, wind regimes, satellite-derived primary production and composition of abundant phytoplankton species. After removing the effects of the most important environmental variables that were correlated with these geographic regions and that likely exert selection (e.g. temperature), we found that differences among Longhurst provinces still accounted for ~20% of picoeukaryotic community variance when using Bray-Curtis and gUniFrac β-diversity indices (PERMANOVA *R^2^*) **[Fig. 2]**. Likewise, Longhurst provinces explained ~20-25% of prokaryotic community variance with all tested β-diversity indices (PERMANOVA *R^2^*) **[Fig. 2]**. The variability in community composition associated to these provinces most likely represent dispersal limitation, even though abiotic or biotic selection exerted by unmeasured variables cannot be ruled out. ß-diversity in picoeukaryotes and prokaryotes displayed positive correlations with distance (i.e. distance decay) predominantly within 1,000 km **(Fig. 5)**, although correlations were weaker in prokaryotes than in picoeukaryotes, being consistent with a higher dispersal limitation in picoeukaryotes than in prokaryotes **(Fig. 1)**.

**Figure 5.**
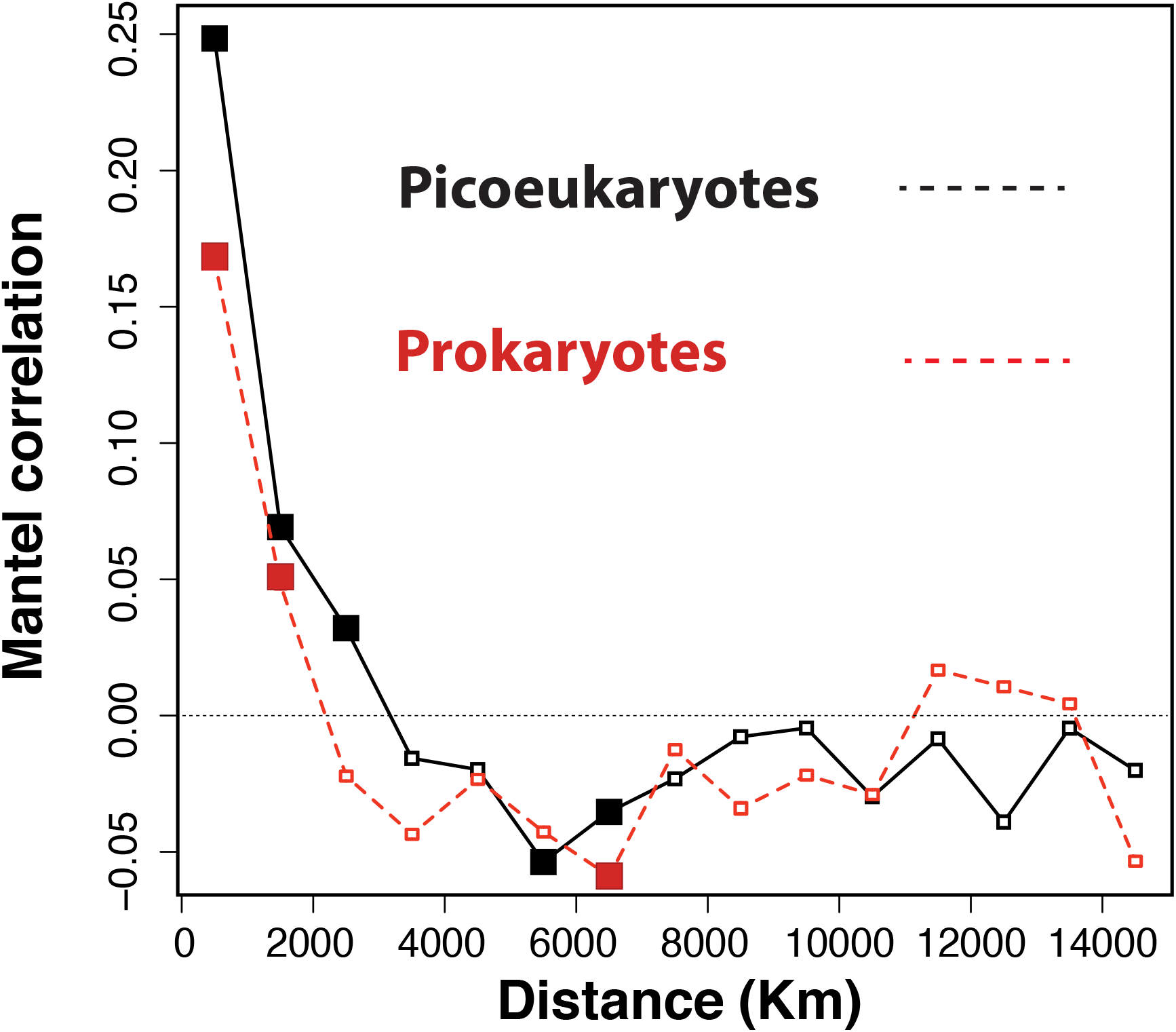
Decrease in community similarity with distance (distance decay). Mantel correlograms between geographic distance and β-diversity featuring distance classes of 1,000 km for both picoeukaryotes and prokaryotes. Coloured squares indicate statistically significant correlations (p<0.05). Note that β-diversity in picoeukaryotes displayed positive correlations with increasing distances up to ~3,000 km, while prokaryotes had positive correlations with distances up to ~2,000 km. Correlations tended to be smaller in prokaryotes than in picoeukaryotes, indicating smaller distance decay in the former compared to the latter.

Selection and dispersal limitation may operate more strongly in geographic areas considered ecological boundaries, for example, due to strong physicochemical change in the seawater, leading to abrupt changes in microbiome composition. We identified a total of 14 communities where sequential β-diversity displayed abrupt changes, with 11 of them coinciding for both picoeukaryotes and prokaryotes **[Fig. 4, Panels A** & **B]**. The Local Contribution to Beta Diversity (LCBD) index (40) **(Fig. 4, Panel D)** indicated that ~22% of both picoeukaryotic and prokaryotic communities (26 stations each, totaling 36 stations) contributed the most to the β-diversity, with 16 communities coinciding for both prokaryotes and picoeukaryotes (p<0.05; **Fig. 4, Panel D, Table S7**). In addition, 8 of the 36 stations featuring a significant LCBD were also identified as zones of abrupt community change in sequential Bray Curtis analyses **(Table S7)**. These zones featuring abrupt community change in both prokaryotes and picoeukaryotes point to selection or dispersal acting simultaneously and strongly upon both life’s domains.

## DISCUSSION

Our assessment of the tropical and sub-tropical sunlit fraction of the global-ocean microbiome during the Malaspina 2010 Circumnavigation Expedition indicated that dispersal limitation is the dominant process structuring picoeukaryotic communities, while selection, dispersal and drift have a balanced importance in structuring prokaryotic communities. These results summarise the general action of ecological processes at the microbiome level and cannot be extended to every single species or group; for example, some picoeukaryotes display cosmopolitan distributions (41). Our results also reflect the action of ecological processes in the tropical and subtropical sunlit ocean. Therefore, considering other zones, such as the polar oceans, could modify the relative importance of these processes. To determine the action of selection, we used the principle indicating that phylogenetically closely related taxa tend to be ecologically similar (and vice versa). This principle was supported by data from prokaryotes and unicellular eukaryotes (42–45). Yet, there can be exceptions (46), and failure to detect selection could inflate the estimates of dispersal limitation in our methodology. We consider that dispersal limitation in picoeukaryotes was not inflated in our analyses, as picoeukaryotes generally displayed more limited spatial distributions than prokaryotes and a higher sequential β-diversity. Drift was pragmatically analysed as the compositional variation between communities that does not differ from random community assembly, thus representing neutral metacommunity dynamics (20, 22, 44). Hence, our quantifications of drift do not reflect the impact of random demography in single communities. Lastly, our estimates of the importance of ecological mechanisms consider taxa with high or moderate abundances, which typically carry biogeographic information (47), thus, our results do not reflect the processes shaping the rare biosphere (48).

Selection, which is known to have an important role in structuring prokaryotic communities (23, 24), explained a higher proportion of community turnover in prokaryotes (~34% of the turnover) than in picoeukaryotes (~17% of the turnover). This modest role of selection in structuring the tropical and subtropical sunlit-ocean microbiome is consistent with the weak environmental gradients characterizing this habitat. In other habitats featuring a high selective pressure, such as Antarctic waterbodies that display a strong salinity gradient, the role of selection in structuring bacteria has been reported to be much higher, accounting for up to ~70% of the community turnover (49). The measured relative importance of selection is also a consequence of the global scale of our survey. For example, in comparatively small marine areas, where dispersal limitation is expected to be low (19), the relative importance of selection could increase. In surface waters of the East China Sea it was found that selection was ~40% more important than dispersal limitation in structuring bacterial communities (50), while in our study, selection and dispersal limitation had a similar importance in structuring prokaryotes. The same study (50) also found that selection was ~40 times more important than dispersal limitation in structuring communities of microbial eukaryotes. In contrast, our global assessment yields dispersal limitation to be ~5 times larger than selection in structuring picoeukaryotic communities.

Heterogeneous selection was more important in structuring picoeukaryotic (~16%) than prokaryotic communities (~9%), while homogeneous selection was more important in prokaryotic (~24%) than in picoeukaryotic communities (~1%). Homogeneous environmental conditions should lead to homogeneous selection, constraining community divergence, while heterogeneous environmental conditions should promote community divergence (18). Our results suggest that prokaryotes and picoeukaryotes in the same marine habitats respond differently to the same environmental conditions. Thus, selection would be preventing community divergence in prokaryotes while promoting it in picoeukaryotes. The fundamental cellular differences between prokaryotes and picoeukaryotes (10, 11) may determine such different responses to the same environmental heterogeneity. For example, comparable environmental heterogeneity could select for a few species featuring wide environmental tolerance or several species, which are adapted to narrow environmental conditions.

Previous studies indicate that temperature is one of the most important variables exerting selection on ocean prokaryotes (51–59). Earlier work (51) reported strong correlations between prokaryotic ocean-microbiome composition and temperature, and weak correlations with nutrients, consistent with our results. Less is known about the effects of temperature on the community structuring of ocean picoeukaryotes, which according to our results are minor. Yet, it is known that temperature affects the distributions of MAST-4, a lineage of ubiquitous marine heterotrophic flagellates (41), suggesting that the effects of temperature on small eukaryotes could be group specific. Interestingly, our results suggest that selection, operating via temperature, affects prokaryotic taxa co-occurrences, having limited effects on picoeukaryotic co-occurences. In prokaryotes, the β-diversity associated to temperature explained by TINA_w_ (~50%) was substantially higher than Bray Curtis (~15%), reflecting the importance of considering co-occurrences, as in TINA_w_, to understand community structure. These results suggest that temperature-driven selection determines the species that can grow in different locations, yet in each site, species relative abundances and presence-absence may vary due to local stochasticity (60, 61).

To what extent dispersal limitation affects the distribution of microbes is a matter of debate (62–65). In surface open-ocean waters, prokaryotes typically display abundances of 10^6^ cells/mL, while picoeukaryotes normally have abundances of 10^3^ cells/mL (66). Due to random dispersal alone, the more abundant prokaryotes are expected to be distributed more thoroughly than the less abundant picoeukaryotes (34), which is consistent with our findings. Furthermore, the absence of taxa from suitable habitats that are separated by large distances is expected to be more pronounced in picoeukaryotes than in prokaryotes. Still, our analyses compare actual species distributions against those that would be expected by chance when considering species abundances, suggesting that species-abundance is not the main reason of dispersal limitation in picoeukaryotes.

Several studies in aquatic unicellular eukaryotes point to restricted dispersal and endemism (64, 67–69), while others indicate the opposite (41, 70–72). This could reflect different dispersal capabilities among small eukaryotic taxa (64, 73) and the generation of dormant cysts in some species, such as in diatoms, dinoflagellates and coccolitophorids (74, 75), which may increase dispersal rates. Cyst formation has not been reported yet for picoeukaryotes (11) and this may partially explain their limited dispersal. Regarding prokaryotes, other studies considering large geographic scales indicated that dispersal limitation has a small influence in the structure of marine communities (51, 52, 76), which is coherent with our results. Dormancy in prokaryotes seems to be more common than in picoeukaryotes (11, 31), and this may allow the former to disperse more thoroughly by reducing their metabolisms when moving through unfavorable habitats (77). In sum, our quantifications of dispersal limitation agree, in general terms, with known trends in both picoeukaryotes and prokaryotes.

The importance of drift or neutral dynamics (20) in structuring microbial communities is also a matter of debate (23, 78). Our results indicate that drift has a modest role in structuring the sunlit-ocean microbiome, being higher in prokaryotic (~31% of the turnover) than in picoeukaryotic communities (~6% of the turnover). Another study also found a higher importance of drift in determining the community structure of bacteria when compared with phytoplankton populating freshwater and brackish habitats (79). In contrast, drift was the prevalent community-structuring mechanism in unicellular eukaryotes populating lakes in a relatively small geographic area that features a strong salinity gradient, having a low importance for the structuring of prokaryotic communities (49). Likely, the relative importance of drift in structuring prokaryotes or unicellular eukaryotes is dependent on the selective strength of specific habitats, the occurrence of adaptive processes (49) or barriers to dispersal.

When geographic distance is correlated with environmental variation, a decrease in community similarity with distance (distance decay) can be the manifestation of both selection and/or dispersal limitation (24). Distance decay has been evidenced in diverse studies focusing on the surface and deep ocean microbiome (52, 80, 81). Yet, different to most previous studies, we have quantified the role of selection and dispersal limitation in structuring the surface ocean microbiome, and we can use this information to interpret the measured distance decay. As our quantifications indicated a strong dispersal limitation in picoeukaryotes, it is likely that this process explains the measured distance decay. In contrast, the distance decay observed in prokaryotes could be the outcome of both selection and dispersal limitation, as both presented comparable structuring roles. The amount of community variance in prokaryotes and picoeukaryotes associated to provincialism (i.e. Longhurst oceanic regions) likely reflects dispersal limitation, since the effects of important environmental variables were removed during the analyses. Interestingly, another study investigating surface marine bacteria along ~12,000 km in the Atlantic Ocean found that provincialism explained an amount of community variance comparable to our results (52). Furthermore, a study of the eukaryotic microbiome in the sunlit global-ocean indicated that provincialism (considered in terms of ocean basins) was one of the most important variables explaining community structure (67). In the light of our findings, we consider that results from the previous study (67) manifest, to a large extent, dispersal limitation.

In the surface ocean, drastic changes in species composition across space may point to strong changes in abiotic selection, as expected to occur across oceanographic fronts (82, 83), or high immigration from the surface or deeper water layers. We identified 14 stations featuring abrupt changes in prokaryotic or picoeukaryotic community composition as well as 36 stations with a “unique” species composition according to the *Local Contribution to Beta Diversity* analysis (40). Several of these stations coincided for both picoeukaryotes and prokaryotes. Some of these areas correspond to nutrient-rich coastal zones (the South African Atlantic coast and the South Australia Bight) or potential upwelling zones, such as the Equatorial Pacific and Atlantic as well as the Costa Rica Dome. This agrees with a scenario including strong selective changes or immigration from deep water layers into the surface, which affects both prokaryotes and picoeukaryotes.

Altogether, our results represent a significant contribution towards understanding the structure of the sunlit-ocean microbiome by connecting patterns to underlying ecological processes (24). Our findings indicate that comprehending the idiosyncrasies of the main components of microbiomes is needed in order to attain a holistic understanding of their structures and ecologies. In particular, our results suggest that the structure of surface-ocean prokaryotic communities could be more susceptible to global warming than that of picoeukaryotic communities. Prokaryotes represent an important fraction of the total microbial biomass in the ocean (84, 85), and they have fundamental roles for ecosystem function (1, 2). Therefore, understanding the specific effects of temperature in their distributions and community metabolism (27) represents an important challenge, which our results contribute to address.

## METHODS

### Sample collection

Surface waters (3 m depth) from a total of 120 globally-distributed stations located in the tropical and sub-tropical global ocean **(Fig. S1)** across ~50,000 km, were sampled from December 2010 to July 2011 as a part of the Malaspina-2010 expedition (33, 86). Water samples were obtained with a 20 L Niskin bottle deployed simultaneously to a CTD profiler that included sensors for conductivity, temperature, oxygen, fluorescence and turbidity. About 12 L of seawater were sequentially filtered through a 20 μm nylon mesh, followed by a 3 μm and 0.2 μm polycarbonate filters of 47 mm diameter (Isopore, Millipore). Only the smallest size-fraction [0.2 −3 μm, here called “picoplankton”; see (5)] was used in downstream analyses. Samples for inorganic nutrients (NO_3_^−^, NO_2_^−^, PO_4_^3-^, SiO_2_) were collected from the Niskin bottles and measured spectrophotometrically using an Alliance Evolution II autoanalyzer (87). Chlorophyll measurements were obtained from Estrada et al. (86). In specific samples nutrient concentrations were estimated using the World Ocean Database (88–91) due to issues with measurements. Since not all environmental parameters were available for all stations, two contextual datasets were generated: *Meta-119*, including 119 stations **(Fig. S1)**, 5 environmental parameters and 5 spatial features and *Meta-57* **(Fig. S4)**, including 57 stations and 17 environmental parameters. See Supplementary Information for further details.

### DNA extraction, 18S- & 16S-rRNA amplicon sequencing and bioinformatic analyses

DNA was extracted using a standard phenol-chloroform protocol (92). Both the 18S and 16S rRNA-genes were amplified from the same DNA extracts. The hypervariable V4 region of the 18S (~380 bp) was amplified with the primers TAReukFWD1 and TAReukREV3 (93), while the hypervariable V4-V5 (~400bp) region of the 16S was amplified with the primers 515F-Y - 926R (94), which target both Bacteria and Archaea. Amplicon libraries were then paired-end sequenced on an *Illumina* MiSeq platform (2×250bp) at the Research and Testing Laboratory facility (Lubbock, TX, USA; http://www.researchandtesting.com/).

Reads were processed following an in-house pipeline (95). Briefly, raw reads were corrected using BayesHammer (96) following Schirmer *et al*. (97). Corrected reads were merged with PEAR (98) and sequences >200bp were quality-checked and de-replicated using USEARCH (99). Operational Taxonomic Units (OTUs) were delineated at 99% similarity using UPARSE V8.1.1756 (100). To obtain OTU abundances, reads were mapped back to OTUs at 99% similarity. Chimera check and removal was performed both *de novo* and using the SILVA reference database (101). After our stringent quality control (see Supplementary Information), a total of 42,505 picoeukaryotic and 10,158 prokaryotic OTUs were obtained. Taxonomic assignment of picoeukaryotic and prokaryotic OTUs was generated by BLASTing (102) representative sequences against different reference databases. Metazoan, Charophyta, nucleomorphs, Chloroplast and mitochondrial OTUs were removed from the OTU tables. See more details in Supplementary Information and **Table S8**. Sequences are publicly available at the European Nucleotide Archive (http://www.ebi.ac.uk/ena; accession numbers PRJEB23913 [18S] & PRJEB25224 [16S]).

In specific analyses, we considered publicly-available data from the TARA Oceans expedition (36). We selected data from surface communities only, including 41 samples (40 stations) for pico-nano eukaryotes (0.22-3 μm [1 sample] and 0.8-5 μm [40 samples]; 18S-V9 rDNA amplicon data) (67) as well as 63 stations for prokaryotes [picoplankton, 0.22-3 μm (45 samples) and 0.22-1.6 μm (18 samples); 16S rDNA, miTags] (51).

### General analyses and phylogenetic inferences

Both picoeukaryotic and prokaryotic datasets were sub-sampled to 4,060 reads per sample using *rrarefy* in *Vegan* (103), resulting in sub-sampled tables containing 18,881 picoeukaryotic and 7,025 prokaryotic OTUs. All OTUs with mean relative abundances above 0.1% and below 0.001% were defined as regionally abundant or rare respectively (104). Phylogenetic trees were constructed for both the 16S and 18S datasets using OTU-representative sequences. Reads were aligned against an aligned SILVA template using *mothur* (105). Afterwards, poorly aligned regions or sequences were removed using *Trimal* (106). A phylogenetic tree was inferred using *FastTree* v2.1.9 (107). Most analyses were performed in the R statistical environment (108) using *adespatial* (109), *APE* (110), *ggplot2* (111), *gUniFrac* (112), *Maps* (113), *Mapplots* (114), *Picante* (115) and *Vegan* (103). See further details in Supplementary Information.

### Quantification of selection, dispersal and drift

Selection, dispersal and drift were quantified using the approach proposed by Stegen et al. (22). This methodology consists of two main steps: the first uses phylogenetic turnover to infer the action of selection and the second uses OTU turnover to infer the action of dispersal and drift. Phylogenetic turnover was measured by calculating the abundance-weighted β-mean nearest taxon distance (βMNTD), which quantifies the mean phylogenetic distances between the evolutionary closest OTUs in two communities. βMNTD values can be larger, smaller or equal to the values expected when selection is not affecting community turnover (that is, expected under a random distribution [null model]). βMNTD values higher than expected by chance indicate that communities are under heterogeneous selection (18). In contrast, βMNTD values which are lower than expected by chance indicate that communities are experiencing homogeneous selection. Null models were constructed using 999 randomizations as in Stegen et al. (22). Differences between the observed βMNTD and the mean of the null distribution are denoted as β-Nearest Taxon Index (βNTI), with | βNTI | > 2 being considered as significant departures from random phylogenetic turnover, pointing to the action of selection.

The second step of this method calculates whether the observed β-diversity, based on OTU turnover, could be generated by drift (i.e. chance) or other processes. For this, we calculated the Raup-Crick metric (116) using Bray-Curtis dissimilarities [hereafter RC_bray_], following Stegen et al. (22). RC_bray_ compares the measured β-diversity against the β-diversity that would be obtained if drift was driving community turnover (that is, under random community assembly). The randomization was run 9,999 times and only OTUs with >1,000 reads over the entire dataset were considered in order to prevent any bias due to potential under sampling. RC_bray_ values between −0.95 and +0.95 point to a community assembly governed by drift. On the contrary, RC_bray_ values > +0.95 or < −0.95 indicate that community turnover is driven by dispersal limitation or homogenizing dispersal respectively (116). The previous framework was applied as following: First, we determined the proportion of community pairwise comparisons displaying a |βNTI| > 2, which points to the action of selection. Subsequently, for the pairwise comparisons that did not indicate the action of selection, we calculated the proportion of total comparisons that could be assigned to dispersal limitation, homogenizing dispersal or drift according to their RC_bray_ values. See further details in Supplementary Information.

### Estimation of interaction-adjusted indices

Taxa INteraction-Adjusted (TINA) and Phylogenetic INteraction Adjusted (PINA) indices were estimated following Schmidt et al. (35) TINA is based on taxa co-occurrences while PINA considers phylogenetic similarities (35). In particular, TINA quantifies β-diversity as the average interaction strength between all taxa in different samples. Thus, communities which are identical or include taxa which are perfectly associated will give TINA values of 1. On the other hand, TINA values will approach 0.5 in communities sharing no taxa or having neutral associations, and approach 0 if taxa display high avoidance. Dissimilarity matrices were generated as 1-TINA. Full picoeukaryotic and prokaryotic subsampled OTU tables were used to calculate the abundance-weighted TINA_w_ and PINA_w_.

### Associations between taxa and environmental parameters

We analysed whether OTUs had differential associations with environmental parameters as well as between themselves using different algorithms. Firstly, we used the Maximum Information Coefficient (MIC) which captures diverse relationships (including non-linear and non-functional) between two pairs of variables (38). The Malaspina dataset consisted of 119 stations and 17 environmental parameters (See Supplementary Information for extra details). In the TARA Oceans dataset, prokaryotes were analysed across 63 surface stations (including 8 environmental parameters), while microbial eukaryotes were analysed across 40 surface stations (including 6 environmental parameters) [see Supplementary Information]. In both datasets, MIC analyses were run using CV=0.5, B=0.6, and statistically significant relationships with MIC ≥0.4 (Malaspina) or MIC ≥0.5 (TARA) were considered; significance was assessed using precomputed p-values (38). Non-linear associations were defined as MIC-ρ^2^ >0.2 (38). Secondly, we constructed association networks with the Malaspina dataset considering OTUs with >100 reads using *SparCC* (117) as implemented in *FastSpar* (118). To determine correlations, *FastSpar* was run with 1,000 iterations, including 1,000 bootstraps to infer p-values. We used OTU associations with absolute correlation scores >0.3 and p<0.01. Networks were visualized with *Cytoscape* (119) and their properties determined using *igraph* (120).

## ACKNOWLEDGEMENTS

We thank all scientists and crews from the Malaspina-2010 expedition. RL was supported by a Ramón y Cajal fellowship (RYC-2013-12554, MINECO, Spain). IMD was supported by an ITN-SINGEK fellowship (ESR2-EU-H2020-MSCA-ITN-2015, Grant Agreement 675752) and CRG by a Juan de la Cierva (IJCI-2015-23505, MINECO, Spain) fellowship. This work was supported by the projects Malaspina-2010 Expedition (CSD2008-00077, MINECO, Spain), INTERACTOMICS (CTM2015-69936-P, MINECO, Spain), REMEI (CTM2015-70340-P, MINECO, Spain) and MicroEcoSystems (240904, RCN, Norway). Bioinformatics analyses were performed at the MARBITS platform of the Institut de Ciències del Mar (ICM; http://marbits.icm.csic.es) as well as in MareNostrum (Barcelona Supercomputing Center) via grants obtained from the Spanish Network of Supercomputing (RES) to RL.

